# Lateralized expression of cortical perineuronal nets during maternal experience is dependent on MECP2

**DOI:** 10.1101/787267

**Authors:** Billy Y.B. Lau, Dana E. Layo, Brett Emery, Matthew Everett, Anushree Kumar, Parker Stevenson, Kristopher G. Reynolds, Andrew Cherosky, Sarah-Anne H. Bowyer, Sarah Roth, Delaney G. Fisher, Rachel P. McCord, Keerthi Krishnan

## Abstract

Cortical neuronal circuits along the sensorimotor pathways are shaped by experience during critical periods of heightened plasticity in early postnatal development. After closure of critical periods, measured histologically by the formation and maintenance of extracellular matrix structures called perineuronal nets (PNNs), the adult mouse brain exhibits restricted plasticity and maturity. Mature PNNs are typically considered to be stable structures that restrict synaptic plasticity on cortical parvalbumin+ GABAergic neurons. Changes in environment (i.e. novel behavioral training) or social contexts (i.e. motherhood) are known to elicit synaptic plasticity in relevant neural circuitry. However, little is known about concomitant changes in the PNNs surrounding the cortical parvalbumin+ GABAergic neurons. Here, we show novel changes in PNN density in the primary somatosensory cortex (SS1) of adult female mice after maternal experience, using systematic microscopy analysis of a whole brain region. On average, PNNs were increased in the right barrel field and decreased in the left forelimb regions. Individual mice had left hemisphere dominance in PNN density. Using adult female mice deficient in methyl-CpG-binding protein 2 (MECP2), an epigenetic regulator involved in regulating experience-dependent plasticity, we found that MECP2 is critical for this precise and dynamic expression of PNN. Adult naïve *Mecp2*-heterozygous females (Het) had increased PNN density in specific subregions in both hemispheres before maternal experience. The laterality in PNN expression seen in naïve Het was lost after maternal experience, suggesting possible intact mechanisms for plasticity. Together, our results identify subregion and hemisphere-specific alterations in PNN expression in adult females, suggesting extracellular matrix plasticity as a possible neurobiological mechanism for adult behaviors in rodents.

## Introduction

Perineuronal nets (PNNs) are specialized extracellular matrix structures that can act as physical barriers or modulators of plasticity, restrict axon regeneration, and form molecular brakes that actively control synaptic maturation and the function of cortical parvalbumin+ (PV+) GABAergic interneurons that drive gamma oscillations (Bartos et al., 2002; Begum and Sng, 2017; Bernard and Prochiantz, 2016; Carstens et al., 2016; Deepa et al., 2002; Dityatev et al., 2007; Donato et al., 2013; Durand et al., 2012; Frischknecht et al., 2009; Gundelfinger et al., 2010; Hartig et al., 1992; Hou et al., 2017; Kalemaki et al., 2018; Kosaka and Heizmann, 1989; Krishnan et al., 2015, 2017; Nakagawa et al., 1986; Orlando et al., 2012; Pizzorusso et al., 2002, 2006; Sigal et al., 2019; Sugiyama et al., 2009; Suttkus et al., 2012; Ueno et al., 2018; Vo et al., 2013; de Winter et al., 2016; Ye and Miao, 2013). In rodents, mature PNNs in the adult cortex are thought to be stable structures, inhibitory to plasticity, and perhaps play roles in long term memory such as “engrams” (Carstens et al., 2016; Gogolla et al., 2009; Thompson et al., 2018). However, most of these observations are based on postnatal cortical development (when typical connections in neural circuitry are still forming) and models for neurobiological disorders (where neural circuitry development and function have gone awry). Currently, it is unclear if changes in adult PNN expression occur under normal conditions and behavioral contexts.

PNNs are composed of chondroitin sulfate proteoglycans, hyaluronan glycosaminoglycan chains, link proteins and tenascin-R (Bignami et al., 1992; Carulli et al., 2007; Kwok et al., 2010; Miyata and Kitagawa, 2017). Wisteria floribunda agglutinin (WFA) is commonly used as a marker for PNNs in the cortex and other brain regions (Brückner et al., 1996; Hartig et al., 1992). WFA specifically binds to N-acetyl galactosamine found on most chondroitin sulfate side chains of chondroitin sulfate proteoglycans. In rodents, WFA-labeled PNNs are localized predominantly around soma and proximal dendrites of parvalbumin+ (PV+) GABAergic interneurons of the mature cortex. They interdigitate with synaptic contacts on cortical PV+ GABAergic neurons and regulate experience-dependent synaptic plasticity in the cortex, hippocampus and amygdala (Cattaud et al., 2018; Gogolla et al., 2009; Krishnan et al., 2015, 2017; Miyata et al., 2012; Murthy et al., 2019; Sigal et al., 2019).

In the human brain, decreased numbers of PNNs are associated with pathological conditions such as decreased memory and motor agility (Brückner et al., 2008; Morawski et al., 2004). Mouse models for varying neurological disorders show abnormal/atypical expression of PNNs which, when removed, can greatly improve the associated pathology or behavioral readouts in these models (Berretta et al., 2015; Krishnan, Lau et al., 2017; Pizzo et al., 2016; Reinhard et al., 2019). We have previously shown that precocious or atypical expression of PNNs caused sensory processing deficits in developing male or adult female mouse models for Rett Syndrome, respectively (Krishnan et al., 2015, 2017). Rett Syndrome is a neuropsychiatric disorder predominantly caused by mutations in the X-linked gene, methyl CpG-binding protein 2 (MECP2) (Amir et al., 1999). MECP2 regulates neuronal chromatin architecture and gene transcription in response to neural activity and experience during postnatal life (Becker et al., 2016; Chahrour et al., 2008; Ebert et al., 2013; Skene et al., 2010; Zhou et al., 2006). The known cellular function of MECP2 and the characteristic timing of disease progression led us to hypothesize that MECP2 regulates experience-dependent plasticity in specific neural circuits during windows of enhanced sensory and social experience throughout life; disruptions in timing of these plasticity mechanisms results in atypical responses in behavior. We previously tested this hypothesis using a pup retrieval task in the alloparental care paradigm (Krishnan, Lau et al., 2017).

Parenting is an ethologically relevant social behavior consisting of stereotypic components involving the care and nourishment of young. First-time dams seek and gather wandering/scattered pups back to the nest (pup retrieval), an essential aspect of maternal care. Pup retrieval involves processing of primary sensory cues (auditory, tactile, olfactory) to direct efficient searching and gathering of pups with goal-directed movements back to the nest (Beach and Jaynes, 1956; Lonstein et al., 2015; Stern, 1996). Virgin female mice (naïve) with no previous maternal experience can execute efficient pup retrieval after co-housing (‘surrogates’ or ‘Sur’) with a first-time mother and her pups (Cohen et al., 2011a). This assay allows for interrogation of adult experience-dependent plasticity mechanisms as well as likely non-hormonal, epigenetic mechanisms involving the sensory and motor neural circuits (Stolzenberg and Champagne, 2016). The role of the auditory cortex in pup retrieval and maternal experience is a topic of investigation in many labs (Cohen et al., 2011a; Galindo-Leon et al., 2009; Marlin et al., 2015; Stiebler et al., 1997). They continue to contribute to the understanding of how relevant sensory cues are processed in the maternal brain.

Previously, we found that atypical increase in PNN expression in the auditory cortex of *Mecp2*-heterozygous females caused inefficient pup retrieval (Krishnan, Lau et al., 2017). However, we did not find discernable changes in PNN density in the wild type auditory cortex after successful completion of the pup retrieval task. On the one hand, as PNNs are considered barriers to plasticity, we anticipated reduction in PNN expression in wild type that could facilitate efficient retrieval. On the other hand, there are no known reports of reduction in PNN expression in normal adult wild type brains. Here, we seek to answer if mature PNNs are maintained as stable structures or undergo dynamic expression changes in the primary somatosensory cortex (SS1) of adult female mice after maternal experience. We focused on the SS1 due to its known roles in tactile sensation, which is also important for efficient pup retrieval (Brecht, 2007; Feldman and Brecht, 2005; Kenyon et al., 1981; Morgan et al., 1992). By using WFA as a marker and whole-brain analysis of SS1, we find that mature PNNs in adult SS1 1) are differentially expressed in a hemisphere- and subregion-specific manner, 2) show dynamic expression changes after maternal experience, and 3) are influenced by MECP2, a DNA methylation reader/epigenetic regulator of chromatin and gene expression.

## Materials and Methods

### Animals

All experiments were performed in adult female mice (10-12 weeks old) that were maintained on a 12hr light-dark cycle (lights on 07:00 h) and received food ad libitum. Genotypes used were CBA/CaJ, *MeCP2*^*Het*^ (C57BL/6 background; B6.129P2(C)-*Mecp2*^*tm1.1Bird*^/J) and *MeCP2*^*WT*^-siblings (Guy et al., 2001). All procedures were conducted in accordance with the National Institutes of Health’s Guide for the Care and Use of Laboratory Animals and approved by the University of Tennessee-Knoxville Institutional Animal Care and Use Committee.

### Pup retrieval behavior

Pup retrieval behavior was performed as previously described (Krishnan, Lau et al., 2017) (Figure 1A). Briefly, we housed two virgin female mice (one *MeCP2*^*WT*^ and one *MeCP2*^*Het*^ termed ‘naïves’, NW and NH respectively) with a first-time pregnant CBA/CaJ female beginning 1-5 days before birth. Upon cohousing, the two naïve mice are now termed ‘surrogates’ (SW for surrogate *Mecp2*^*WT*^ and SH for surrogate *Mecp2*^*Het*^). Pup retrieval behavior started on the day the pups were born (postnatal day 0; D0) as follows for each adult surrogate mouse: 1) (habituation phase) one adult mouse was habituated with 3-5 pups in the home cage for 5 minutes, 2) (isolation phase) pups were removed from the cage for 2 minutes, and 3) (retrieval phase) pups were returned to home cage, one placed at each corner and at the center (the nest was left empty if there were fewer than 5 pups). Each adult female had maximum of 10 minutes to gather the pups to the original nest. After testing, all animals and pups were returned to the home cage. The same procedure was performed again daily to D5. All behaviors were performed in the dark, during the light cycle (between 9:00 AM and 6:00 PM) and were video recorded.

**Figure 1:**
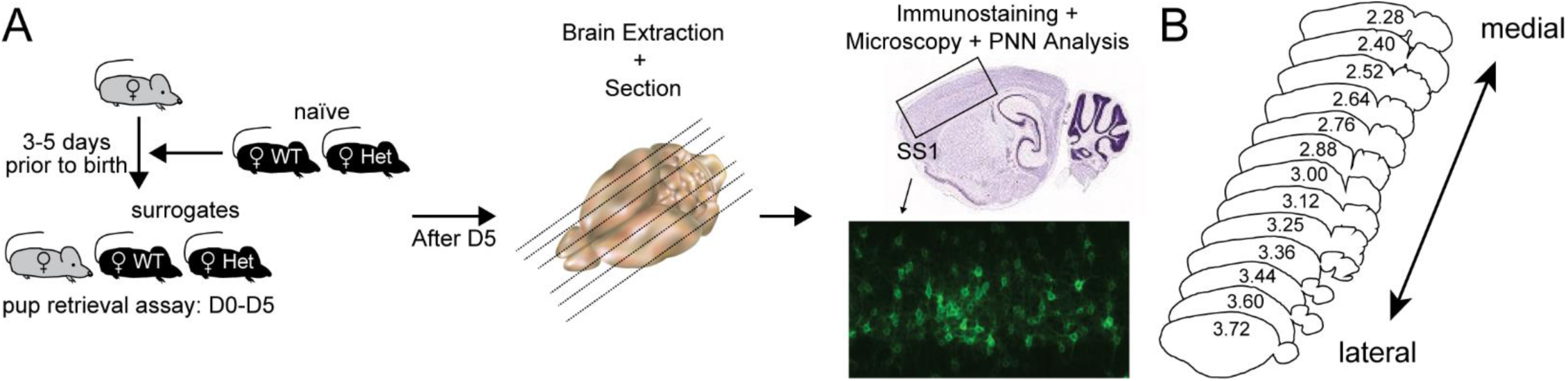
Schema representing behavioral and histology pipeline. **(A)** (Left) Alloparental behavioral model in mice. Pregnant CBA/CaJ female (grey mouse) is cohoused with adult female naïve WT and naïve Het littermate controls, which changes their status to surrogates, 3-5 days before birth of pups. Once pups are born, pup retrieval assay is performed with the surrogates from day 0 (D0) to 5 (D5). (Center) After behavioral experiments on D5, surrogate mice and age-matched naïve counterparts are perfused, their brains extracted, and sectioned as a single cohort. (Right) Standard immunostaining and imaging with epifluorescent slide scanner are performed, to image and analyze PNNs, as elaborated in Methods. **(B)** Schema of mouse brain sections cut in sagittal orientation, depicting all medial (2.28 mm, corresponding to map number 120) to lateral (3.72 mm, corresponding to map number 132) regions of SS1 analyzed for this study. Coordinate maps are based on Paxinos and Franklin’s mouse brain atlas, 4^th^ edition.

### Immunohistochemistry

Immediately after the behavioral trial on D5, surrogate mice as well as a set of corresponding naïve *MeCP2*^*WT*^ and *MeCP2*^*Het*^ mice were perfused with 4% paraformaldehyde/PBS, and brains were extracted and post-fixed overnight at 4°C. Brains were then treated with 30% sucrose/PBS overnight at room temperature (RT) and sectioned sagittally using a freezing microtome at 70μm. Free-floating brain sections were immunostained at RT as previously described in Krishnan, Lau et al., 2017, with a few modifications. Briefly, sections were blocked in 10% normal goat serum and 0.5% Triton-X for 3 hours, then incubated with biotin-conjugated Wisteria Floribunda Agglutinin Lectin (labels PNNs; 1:500; Sigma-Aldrich) overnight at RT in a 5% normal goat serum/0.25% Triton-X solution. Then, sections were incubated for 4 hours with AlexaFluor-488 secondary antibody (1:1000; Invitrogen) in a 5% normal goat serum/0.25% Triton-X solution. Finally, sections were counterstained with the nuclear marker, DAPI (1:1000) for 5 minutes, and mounted in Fluoromount-G (Southern Biotech).

### Image Acquisition and analysis

To analyze PNNs, 10X single-plane PNN images of the entire primary somatosensory cortex from each brain slice were acquired on a motorized stage, epifluorescent microscope (Keyence BZ-X710; Keyence Corp., NJ, U.S.A.) and stitched using BZ-X Analyzer (Keyence) (Figure 1B). Our initial observation of PNN intensity showed that SH had the most intense fluorescent signal in SS1. Thus, imaging settings were established based on SH within each cohort of mice, in order to minimize over-exposure. The light exposure time for fluorescent signal acquisition was identified by finding the exposure time where a saturated pixel first appears within frame, then decreasing the exposure time by 1 unit, according to software specifications. This exposure time determination method was applied to all brain sections of SH in each cohort and the mean exposure time was calculated and used for final image acquisition within each cohort.

For image analysis, each stitched image was opened in ImageJ (Schneider et al., 2012). Then, SS1 and somatosensory subregion areas were 1) mapped by overlaying templates from Paxinos and Franklin’s “The Mouse Brain, 4^th^ edition” (Paxinos and Franklin, 2013), 2) outlined and 3) measured using the functions in ImageJ. To count high-intensity mature PNNs, the ‘Contrast’ setting from the browser was set to the far right to threshold weaker signals. The remaining signals were manually quantified under the classification that a ‘mature’ PNN is at least 80% of its original shape (before contrast adjustment). All statistical analyses and graphs were generated using GraphPad Prism.

### Principal Components Analysis (PCA) on PNN density

We used PNN densities (counts per area) for the entire SS1 across all individuals in the five cohorts. If multiple sections per map number were present, values were averaged across sections to give a single density. In the first PCA analysis, to determine whether the PNN patterns segregated primarily by cohort or (genotype and experience), we kept the data for each individual animal separate and averaged PNN densities across every set of 2 adjacent map regions. Because the scale of raw numbers varied so widely between individual animals, we next median normalized PNN densities within each animal. To do this, we calculate the median PNN density across all map numbers for each individual. Then, each individual PNN density for each map number of that individual was divided by (‘normalized by’) this median. Finally, these median normalized densities were averaged across all individuals in the same condition. This method of normalization accounts for the variability in the median across cohorts within the same condition. Some regions for some individuals had no data, and, because PCA does not tolerate missing data, we imputed these missing values by using the average median normalized values for that region from animals in other cohorts in the same condition. By running PCA in R, we obtained a weight for each map number showing the highest variance patterns in PNN densities across the individual animals. We then calculated the projection of each animal onto principle components 1 and 2. K-means clustering was used to calculate the two most evident clusters in this PCA space.

In the second PCA analysis, to determine the major PNN density patterns that distinguish genotypes and experience conditions, we left data for each map number separate, but averaged the median normalized PNN densities across the 5 cohorts for each condition. We then performed PCA to determine the major patterns that distinguish these conditions.

## Results

### PNN density changes across somatosensory cortical maps

As somatosensation during pup retrieval primarily involves facial/snout areas in surrogates (non-lactating adult females) (Lonstein and Stern, 1997; Morgan et al., 1992; Stern, 1996), we present data collected from the somatosensory cortical regions involved in processing tactile stimuli in the alloparental care paradigm (Figure 1A). According to Paxinos and Franklin’s atlas (4^th^ edition), there are eight different anatomical somatosensory cortical subregions (S1, S1BF, S1DZ, S1FL, S1J, S1ULp, S1Tr and S1HL), here collectively called primary somatosensory cortex (SS1) (Paxinos and Franklin, 2013). To determine how PNN expression changes across the different SS1 subregions, we took a systematic approach covering the whole SS1, rather than the standard approach of analyzing “representative sample sections” (Figure 1B). We analyzed 40-60 sagittal brain sections (at 70um each, both left and right hemispheres) per animal, in five biological replicates across four conditions [naïve WT (NW), naïve Het (NH), surrogate WT (SW), surrogate Het (SH)] (Figure 1A), as qualitative regional differences in PNN density were observed in pilot studies in our lab. The SS1 subregions are represented by map numbers 120-132, corresponding to lateral coordinates 2.28 mm (medial region) to 3.72 mm (lateral region), which encompass ∼1.5 mm of one mouse brain hemisphere.

Across the five cohorts, PNN density (as measured by high-intensity PNN counts over area) was not significantly different between NW and SW across the whole SS1 (Figure 2D). However, we noticed a wide range distribution of PNN density across cohorts, ranging from 1 to 300. In order to determine the source of such variability, we parsed the data according to hemispheres (Figure 2E), across the lateral-medial axis (Figure 3) and by subregions (Figure 4). No significant difference was observed in PNN density between the left and right hemispheres of NW; however, the right hemisphere of SW (SW-R) had increased PNN density compared to the left hemisphere (SW-L) (Figure 2B, C, E), suggesting that maternal experience contributes to increased PNN density in the right hemisphere of SS1, possibly to consolidate new tactile information related to the pups and/or the mother.

**Figure 2:**
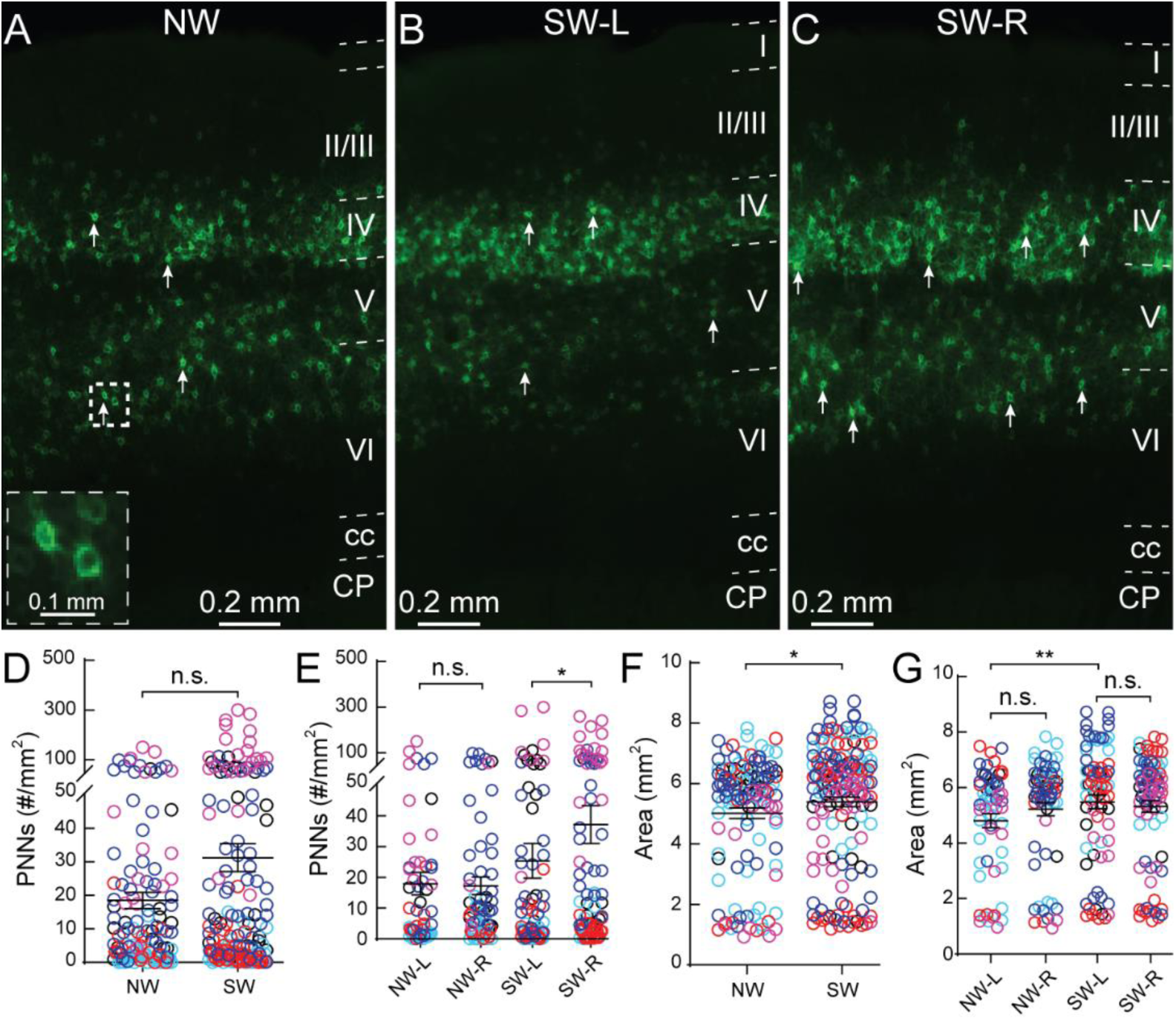
*Mecp2*^*WT*^ (*Wild type*) mice exhibit hemisphere specific increase in PNN density in the primary somatosensory cortex (SS1) after maternal experience. (**A-C**) Representative epifluorescent images of PNN expression in SS1 of naïve WT (NW) (A) as well as left (L) (B) and right (R) (C) hemispheres of surrogate WT (SW). Layers 1 through 6 are outlined. CP = caudate putamen and cc = corpus callosum. Arrows indicate examples of high-intensity PNNs analyzed for the study. Inset in A shows magnified PNN structures in the box. **(D)** Combined hemisphere analysis of the density of high-intensity PNNs was not significantly different between NW and SW (NW: n = 123 images; SW: n = 162; *Mann-Whitney test, p* > *0.05*). **(E)** Separate analysis of left and right hemisphere revealed a significant increase of high-intensity PNNs in the right hemisphere of SW (SW-R; n = 81 images) compared to the left hemisphere of SW (SW-L; n = 81 images) (*Kruskal-Wallis followed by Dunn’s test, *p* < *0.05*), while no significant difference was observed between hemispheres of NW (NW-L: n = 59 images; NW-R: n = 64 images; *Mann-Whitney test, p* > *0.05*). (**F-G**) Size of SS1 was overall significantly larger in SW compared to NW (F) (NW: n = 123 images; SW: n = 162 images; *Mann-Whitney test, *p* < *0.05*). Enlargement of SS1 in SW occurred in the left hemisphere (G) (NW-L: n = 59 images, NW-R: n = 64 images; SW-L: n = 81 images; SW-R: n = 81 images; *Kruskal-Wallis followed by Dunn’s test, **p* < *0.01*). For D-G, n.s. = not significant. Different colors represent each of the five cohorts. Each open circle represents PNN density in an individual brain section.

**Figure 3:**
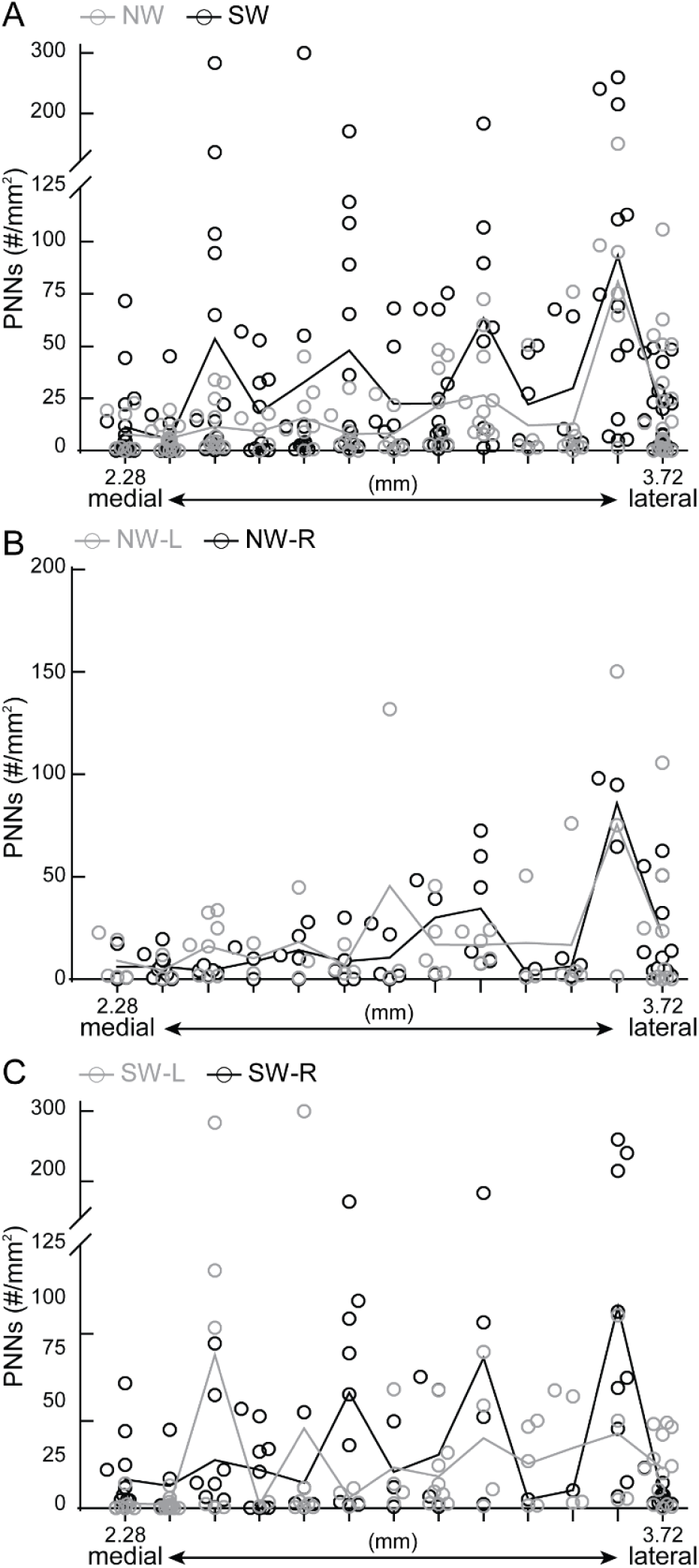
Dynamic changes in PNN density across medial/lateral axis before and after maternal experience. **(A)** Distribution of combined PNN density from left and right hemispheres of SS1 revealed a gradual increase of PNN density from medial (starting at 2.28 mm) to lateral (ends at 3.72 mm) SS1 in NW. After maternal experience, SW exhibited an increased PNN density throughout SS1. N = 5 – 24 images per region, 5 mice per condition. **(B)** In NW, the distribution of PNN density was similar between left and right hemispheres throughout SS1. **(C)** SW exhibited dynamic changes in PNN density between left and right hemispheres as well as throughout medial and lateral SS1. For A-C, lines represent the mean values. Each dot represents PNN density in an individual section. 5 mice per condition. For B and C, n = 1 – 13 images per map coordinate.

**Figure 4:**
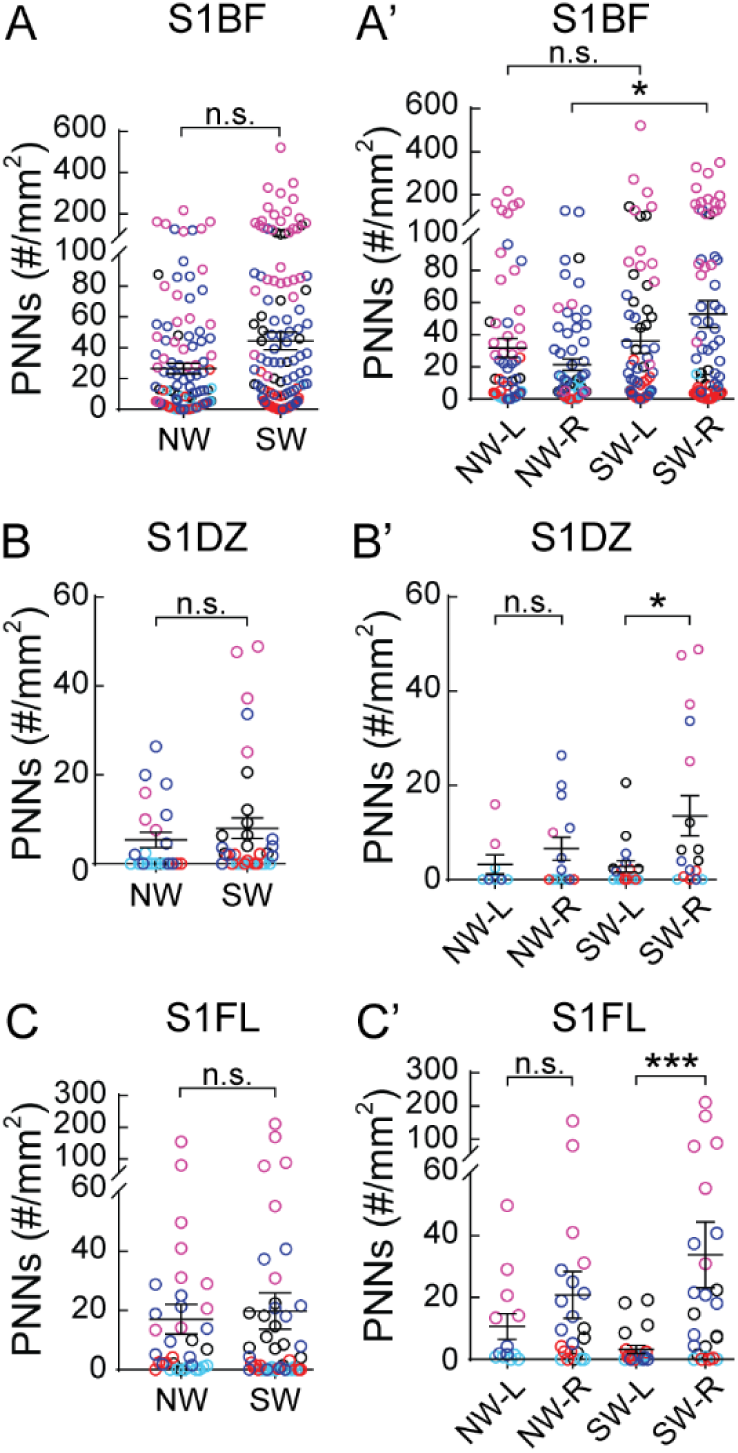
PNN density varies across subregions of SS1. **(A-C)** Analysis of both hemispheres for SS1 barrel field (S1BF, A), dysgranular zone (S1DZ, B) and forelimb (S1FL, C) revealed no significant differences between NW and SW (S1BF: NW - n = 131 images, SW - n = 168 images; S1DZ: NW – n = 22 images, SW – n = 35 images; S1FL: NW – n = 35 images, SW – n = 46 images; 5 mice per condition; *Mann-Whitney test, p* > *0.05, n.s. = not significant*). **(A’-C’)** Analysis of subregional SS1 by hemispheres revealed dynamic changes in PNN expression. In S1BF (**A’**), a significant increase of PNN expression was detected in the right hemisphere of SW (SW-R) compared to NW (NW-R) (NW-L: n = 65 images; NW-R: n = 66 images; SW-L: n = 85 images; SW-R: n = 83 images; 5 mice per condition; *Kruskal-Wallis followed by Dunn’s test, *p* < *0.05*). In S1DZ (**B’**), while no significant difference was observed in PNN density between hemispheres in NW, maternal experience significantly increased PNN density in the right hemisphere (SW-R) compared to the left (SW-L) (NW-L: n = 8 images; NW-R: n = 14 images; SW-L: n = 18 images; SW-R: n = 17 images; 5 mice per condition; *Kruskal-Wallis followed by Dunn’s test, *p* < *0.05*). A similar pattern of PNN plasticity was also detected in S1FL (**C’**), where maternal experience significantly increased PNN density in the right hemisphere than the left of WT (NW-L: n = 13 images; NW-R: n = 22 images; SW-L: n = 21 images; SW-R: n = 25 images; 5 mice per condition; *Kruskal-Wallis followed by Dunn’s test, ***p* < *0.001*).

PNN density is counts over area. Overall, SS1 area of the SW is significantly increased, compared to NW (Figure 2F); however, that increase was not specific to a particular hemisphere within experimental condition (NW-L vs. NW-R; SW-L vs. SW-R) (Figure 2G). After experience, there is a small but significant area increase in left hemisphere (Figure 2G). Thus, the increased PNN density in the right hemisphere of SW is mainly due the PNN expression, and not due to the change in area.

Next, we plotted PNN density in individual sections across the lateral-medial axis (Figure 3). In general, there was a gradual increase in PNN density from medial to lateral SS1, in both NW and SW. However, changes in PNN density after surrogacy experience was seen in more medial sections (Figure 3A). NW had similar PNN density across left (grey) and right (black) hemispheres (Figure 3B), while SW showed hemisphere-specific changes in PNN density across the lateral-medial axis (Figure 3C). These findings indicate that maternal experience can induce changes in the expression of mature PNNs.

### Changes in PNN density are SS1 subregion-specific

Next, we examined if specific subregions of SS1 were particularly plastic for PNN density between NW and SW. We observed no significant differences in PNN density in subregions S1BF, S1DZ and S1FL when aggregating both hemispheres (Figure 4A - C). For S1BF, a region well-studied for whisker activity that contributes to tactile sensation, PNN density increased significantly and specifically in the right hemisphere for SW, compared to NW (Figure 4A’). This result suggests that increased PNN density in the right hemisphere of S1BF could be a potential site for consolidation of tactile sensory information relevant for executing efficient pup retrieval. Contrary to the pattern in S1BF, S1DZ and S1FL regions showed increased PNN density in the right hemisphere, compared to left hemisphere, after surrogacy (Figure 4B’, 4C’). S1DZ has been implicated in proprioceptive functions, such as the movement of joints and stretch of muscle receptors (Chapin and Lin, 1984; Lee and Kim, 2012; Shin Yim et al., 2017; Welker et al., 1984); while S1FL is the sensory representation of the forepaw. Currently, the roles of these regions and PNN contribution in maternal behavior is unclear. Other brain regions (S1ULp, S1J and S1DZ) showed no changes after surrogacy, which further highlights S1BF as a potential site for learning consolidation (Table 1). In analyzing left hemisphere-specific data, there is a decrease in PNN density in S1FL of SW compared to NW (Table 1, columns 5, 7). This is the first report, to our knowledge, of decreases in PNN density in a social behavior context in adult brains. Together, these results suggest that PNN density changes in adult females, in a hemisphere- and subregion-specific manner that is conducive for experience-dependent plasticity.

**Table 1:** Average high-intensity PNN density across subregions and hemispheres of the SS1 before and after maternal behavior experience. NW, NH, SW and SH are the four different conditions. Primary somatosensory cortex subregions: S1BF, S1ULp, S1FL, S1J, S1, and S1DZ. Significant differences are denoted between genotypes by shading (E.g., NW vs NH), between hemispheres of the same condition by **bold** lettering, and between Naïve and Surrogate in the same genotype by bold borders. Each letter pair corresponds to statistically significant differences between two conditions. Numbers correspond to average PNN density with standard error mean across multiple sections. N = 8 – 85 images for hemisphere analysis, 123-168 images for combined hemisphere analysis; 5 mice per condition; *Kruskal-Wallis followed by Dunn’s test*.

| Regions | Whole brain |  |  |  | Left Hemisphere |  |  |  | Right Hemisphere |  |  |  |
| --- | --- | --- | --- | --- | --- | --- | --- | --- | --- | --- | --- | --- |
|  | NW | NH | SW | SH | NW-L | NH-L | SW-L | SH-L | NW-R | NH-R | SW-R | SH-R |
| S1BF | 26.6 ± 3.5<sup>e</sup> | 56.0 ± 6.6<sup>e</sup> | 44.4 ± 5.8 | 47.0 ± 5.4 | 31.7 ± 5.9<sup>f</sup> | 41.0 ± 5.9<sup>f</sup> | 36.1 ± 7.8 | 37.6 ± 5.4 | 21.4 ± 3.6<sup>g,h</sup> | 72.2 ± 11.9<sup>g</sup> | 52.8 ± 8.4<sup>h</sup> | 56.3 ± 9.3 |
| S1ULp | 28.3 ± 4.8<sup>o</sup> | 47.0 ± 8.1<sup>o</sup> | 37.9 ± 5.7 | 52.9 ± 7.6 | 33.4 ± 7.3 | 45.3 ± 8.2 | 29.1 ± 6.7 | 44.2 ± 9.7 | 23.5 ± 6.3 | 49.2 ± 15.2<sup>p</sup> | 38.9 ± 8.0 | 62.5 ± 11.8<sup>p</sup> |
| S1FL | 17.0 ± 5.0 | 20.9 ± 4.0 | 19.8 ± 6.2 | 18.4 ± 3.6 | 10.6 ± 4.2<sup>s</sup> | 20.8 ± 5.0 | 3.2 ± 1.3<sup>s,t,u</sup> | 8.1 ± 2.2<sup>t,v</sup> | 20.8 ± 7.5 | 21.0 ± 6.4 | 33.8 ± 10.7<sup>u</sup> | 30.6 ± 6.6<sup>v</sup> |
| S1J | 2.5 ± 0.8<sup>i</sup> | 4.2 ± 0.9<sup>i</sup> | 6.7 ± 3.6 | 8.4 ± 2.0 | 4.0 ± 1.6<sup>k</sup> | 5.2 ± 1.4<sup>k,l</sup> | 9.0 ± 6.8 | 3.6 ± 1.2<sup>l</sup> | 1.3 ± 0.4<sup>m</sup> | 3.0 ± 0.7<sup>m</sup> | 4.2 ± 1.3<sup>n</sup> | 14.0 ± 3.9<sup>n</sup> |
| S1 | 6.5 ± 0.9<sup>a</sup> | 16.7 ± 2.8<sup>a</sup> | 14.4 ± 3.7<sup>b</sup> | 14.4 ± 2.1<sup>b</sup> | 7.0 ± 1.6 | 11.4 ± 2.5 | 16.7 ± 6.6 | 11.4 ± 2.1 | 6.0 ± 1.0<sup>c</sup> | 22.5 ± 5.0<sup>c</sup> | 12.3 ± 3.2<sup>d</sup> | 17.5 ± 3.6<sup>d</sup> |
| S1DZ | 5.4 ± 1.7 | 9.1 ± 2.6 | 8.0 ± 2.3 | 12.7 ± 3.5 | 3.2 ± 2.0 | 7.4 ± 3.4 | 2.8 ± 1.2<sup>q</sup> | 6.7 ± 4.0<sup>r</sup> | 6.6 ± 2.4 | 10.1 ± 3.8 | 13.4 ± 4.3<sup>q</sup> | 20.6 ± 5.6<sup>r</sup> |

### Appropriate PNN expression in SS1 is dependent on MECP2

Methyl-CpG-binding protein 2 (MECP2) is thought to regulate experience-dependent plasticity mechanisms in an epigenetic manner, in early postnatal development and in adulthood (Cohen et al., 2011b; Dani et al., 2005; Durand et al., 2012; Gabel et al., 2015; Guy et al., 2001; Krishnan et al., 2015, 2017; Lagger et al., 2017; Morello et al., 2018; Muotri et al., 2010; Noutel et al., 2011; Picard and Fagiolini, 2019). We previously tested this hypothesis, using an alloparental care paradigm, and found that *Mecp2*-heterozygous (Het) adult females were inefficient at pup retrieval (Krishnan, Lau et al., 2017). In this study, we identified atypical and transient increases in PNN density in the auditory cortex of surrogate Het (SH), leading to altered responses of PV+ neurons to auditory cues in SH (Lau et al., 2019). Here, we sought to determine if SS1 of SH exhibited similar alterations in PNN density in subregion-specific ways. Comparing naïve Het (NH) to SH, we noticed no significant differences in PNN density in whole SS1 (Figure 5A, similar to WT in Figure 2D), or within hemispheres of SS1 (Figure 5B, unlike WT in Figure 2E), or across the medial-lateral axis (Figure 6). There were no significant changes in SS1 area between NH and SH (Figure 5C; unlike SW in Figure 2F). However, an increased SS1 area in left versus right hemisphere of SH was noted (Figure 5D).

**Figure 5:**
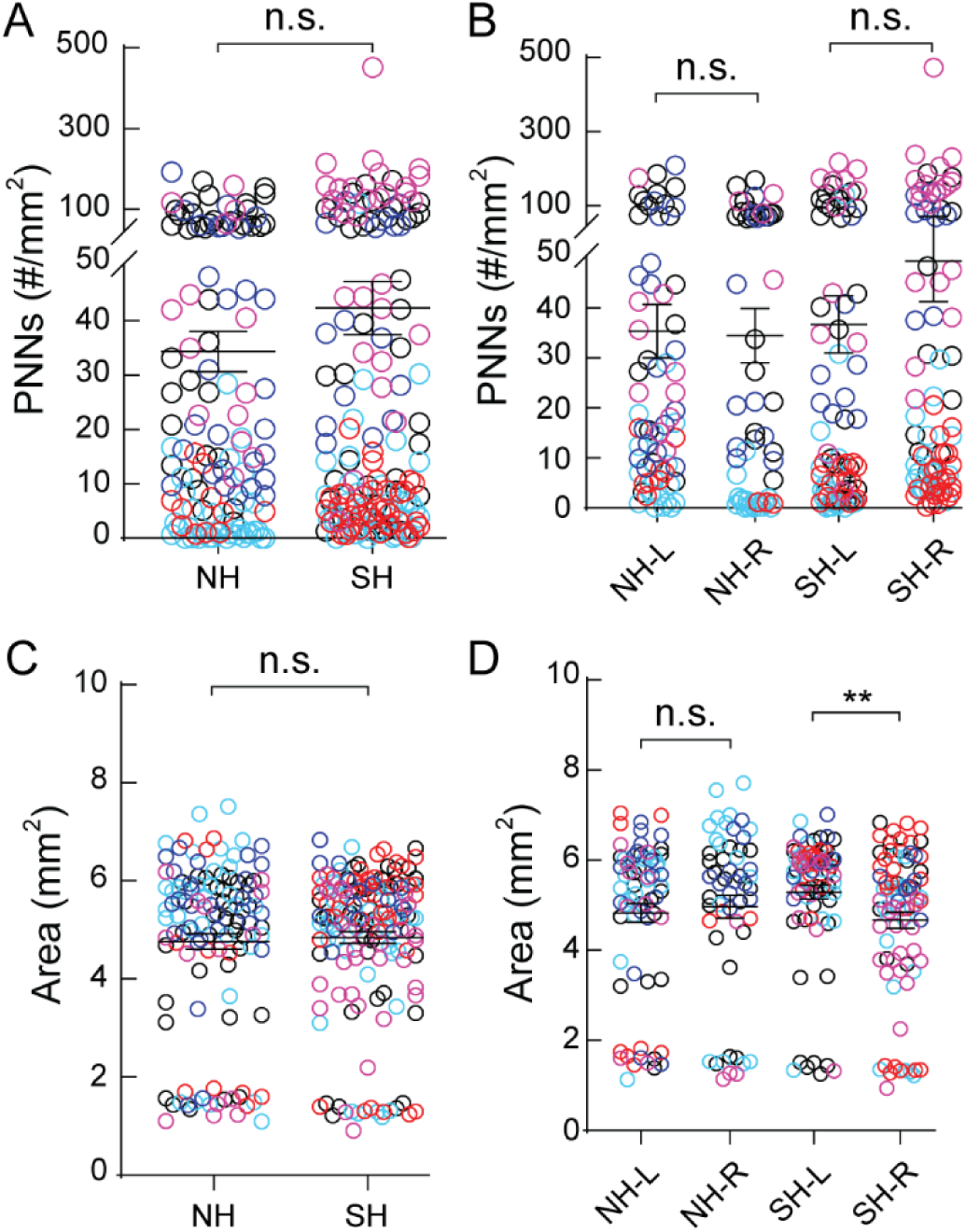
Hemisphere-specific PNN density changes are not conserved in *Mecp2*^*Het*^ after maternal behavior. **(A)** Combined hemispheric analysis of PNN density of SS1 did not reveal significant changes between naïve Het (NH, n = 123 images) and surrogate Het (SH, n = 162 images) (*Mann-Whitney test, p* > *0.05*). **(B)** Analysis of PNN density between hemispheres of SS1 revealed no significant difference between conditions or within hemispheres of naïve or surrogate Het (n = 54 – 82 images; *Kruskal-Wallis followed by Dunn’s test, p* > *0.05*). **(C)** Analysis of SS1 area in both hemispheres revealed no significant changes after maternal learning (NH: n = 123 images; SH: n = 162 images; *Mann-Whitney test, p* > *0.05*). **(D)** Area analysis by hemispheres reveal that left hemisphere of SH (SH-L) was significantly larger than the right hemisphere of SH (SH-R). This hemispheric area bias was absent in NH (n = 54 – 82 images; *Kruskal-Wallis followed by Dunn’s test, **p* < *0.01*). N.s. = not significant. 5 mice per condition.

**Figure 6:**
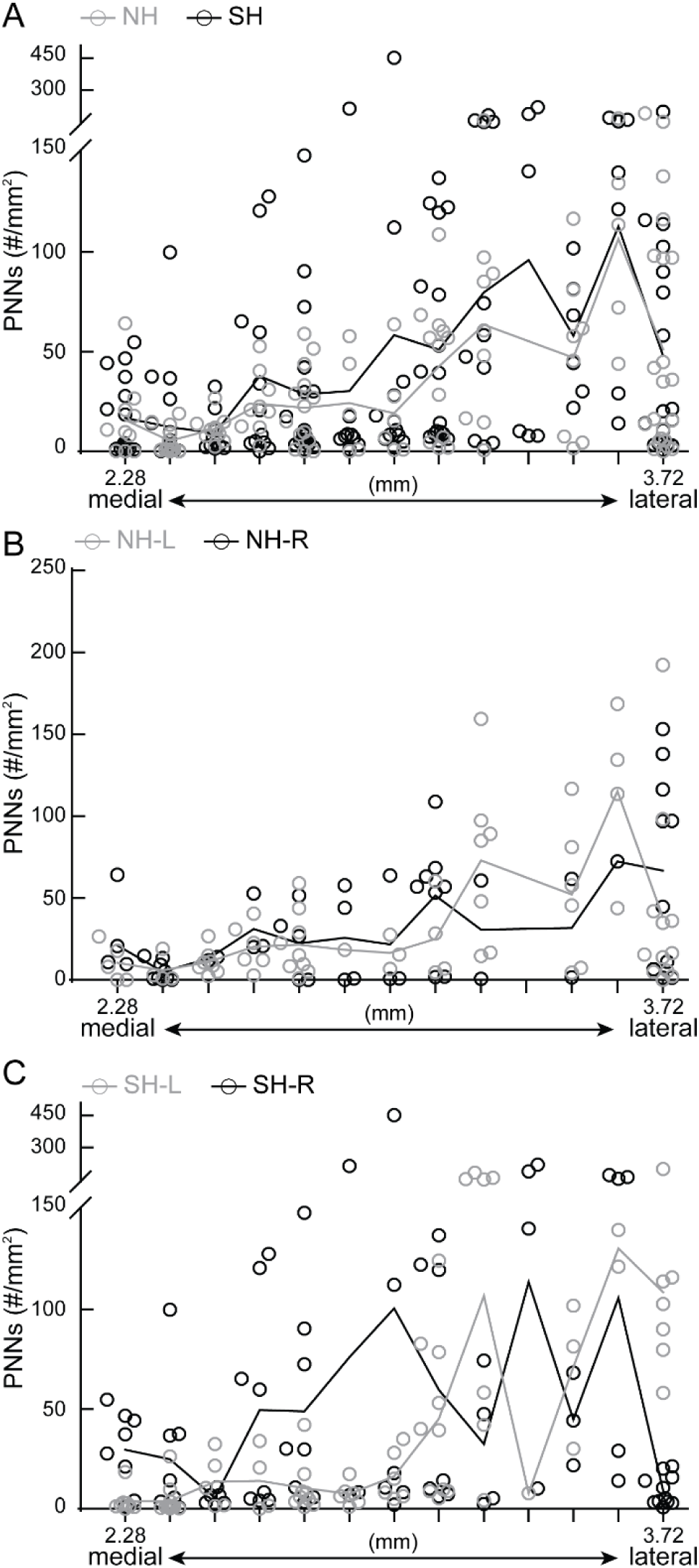
Overall patterns of PNN density across medial/lateral axis is preserved in *Mecp2*^*Het*^. **(A)** Distribution of combined PNN density revealed a gradual increase of PNN density from medial (starting at 2.28 mm) to lateral (ends at 3.72 mm) SS1 in NH. SH exhibited increased PNN density throughout SS1. N = 5 – 22 images per region, 5 mice per condition. **(B)** In NH, distribution of PNN density was similar between left and right hemispheres throughout SS1, with the exception of more lateral regions. **(C)** SH exhibited dynamic changes in PNN density between left and right hemispheres as well as throughout medial and lateral SS1. For A-C, lines represent mean values. Each dot represents PNN density in an individual section. 5 mice per condition. For B and C, n = 1 – 12 images per map coordinate.

When we compared PNN density between genotypes, WT and Het (Table 1), we noticed significant hemisphere-specific and subregion-specific differences. In general, NH had increased PNN density over NW across subregions, and largely no significant differences after surrogacy (SW vs. SH). S1 was an exception, which showed statistical significance, though the mean density was similar. This is likely due to differential distribution of PNN density (Supplemental Figure 1).

**Supplemental Figure 1:**
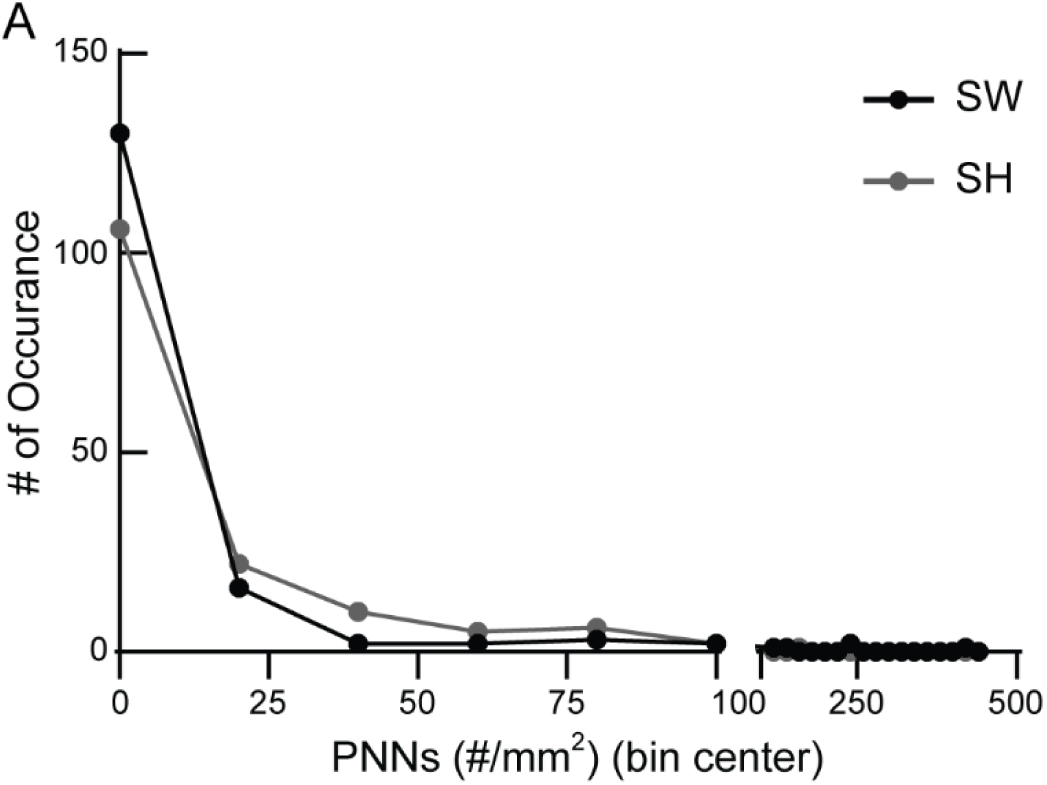
Significant difference in S1 PNN density between SH and SW is explained by differential distribution of data. **(A)** Histogram analysis shows that SW had more occurrences of 0 PNN density, compared to SH, as shown by black dot above grey dot. SH had more occurrences of 20-80 PNN density values than SW, as shown by grey dots above black dots. The average PNN density values of SW (n = 160 images) and SH (n = 152 images) are similar (Table 1) (5 mice per condition), but they are statistically different due to this differential distribution of data.

In the left hemisphere, there were significant differences between conditions, genotypes and subregions. NH had increased PNN density in S1BF and S1J, compared to NW (Table 1, columns 5, 6). SH had increased PNN density in only S1FL, compared to SW (Table 1, columns 7, 8). In the right hemisphere, NH had increased PNN density in S1BF, S1J and S1, compared to NW. SH had increased PNN density in S1J and S1, compared to SW. Surrogacy correlated with increased PNN density in only S1BF of WT (NW vs. SW) (also Fig 2E), while the same increase was not seen in Het (NH vs. SH), with NH-R already exhibiting high PNN density. S1ULp showed increased PNN density in SH-R compared to NH-R (Table 1, columns 10, 12), which was not observed in WT. Taken together, these results suggest that MECP2 regulates dynamic PNN expression, which is then important for appropriate maternal behavior.

Comparing the hemispheres within genotypes, NW and NH did not have significant differences in PNN density in subregions. However, after alloparental experience, PNN density increased in the right hemisphere of S1FL and S1DZ in both SW and SH (Table 1), suggesting preserved common experience-dependent plasticity mechanism activation in these regions within the right hemisphere across genotypes.

### Principal component analysis identifies lateral-medial and hemisphere-specific changes in PNN expression

Due to the increasing number of variables being compared, we chose an unsupervised statistical procedure called principal component analysis (PCA), commonly used in genomics/transcriptomics analysis, to determine if patterns emerge from the PNN density data. PCA takes a set of measurements across samples and identifies the measurements that best capture the variation among the samples. It results in a set of uncorrelated components (called principal components) that each capture an orthogonal aspect of the differences across the samples. As input to PCA, we used PNN densities across individual sections and map numbers (represented as lateral coordinates) across all conditions in the five cohorts. If multiple sections per map number were present, values were averaged across sections to give a single density value.

In the first analysis (Figure 7A, B), we sought to determine whether the PNN patterns segregated primarily by cohort or condition (genotype and experience). We preserved data for each individual brain and performed PCA on PNN densities averaged across every set of 2 adjacent map regions. By examining the projection of each individual onto the first and second principal components, we found that, while there is biological variability between cohorts, the individuals in a given cohort did not cluster separately from one another in this unsupervised analysis (Figure 7A, left), suggesting that technical variability in processing samples across five cohorts is not the primary driver of PNN density differences. This is an important control to assess technical or biological variability in this data. Instead, the first principal component (PC1), which explains 40% of the variation in the data (Figure 7A, right, inset), distinguished the SH PNN density patterns from all others, especially NH (Figure 7A, right). We then examined the weights of each brain region in PC1 to determine which regions are most important for capturing the differences between SH and NH. The weights of each section in PC1 (and PC2) shows a left-right asymmetry and increasing weights for lateral vs. medial sections, confirming our earlier observations (Figure 7B). The findings from our first PCA confirms that variations among mice are resulting from genetic and/or environmental differences and not technical biases. The results further validate our earlier observations of asymmetry in PNN density of left and right hemispheres, augmentation of PNN density in lateral sections, and altered SH PNN density patterns.

**Figure 7:**
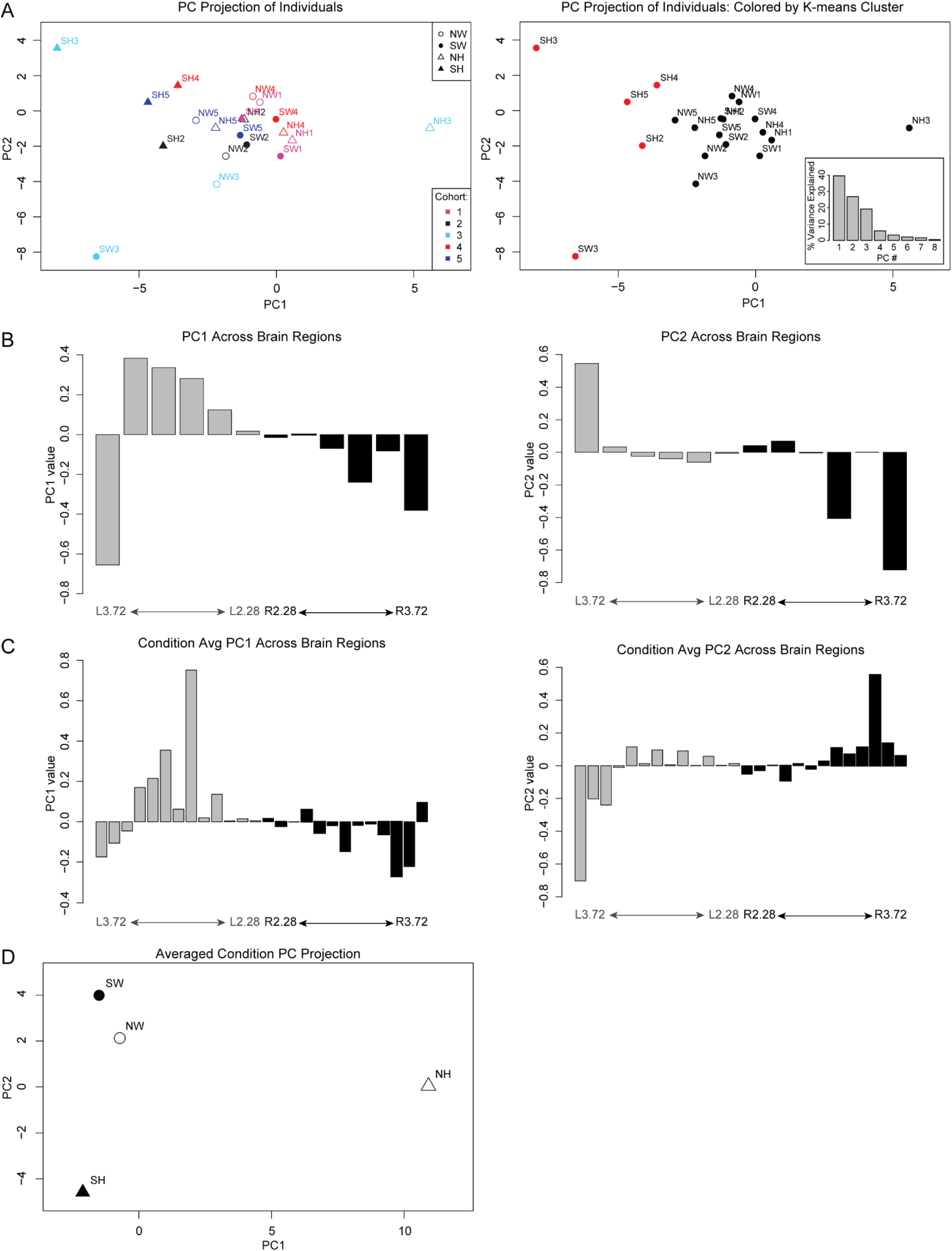
Principal component analysis of PNN expression segregates by conditions, lateral/medial axis and hemispheres. **(A)** The projection of each individual brain onto PC1 and 2. (Left) Individuals are colored by cohort with symbol shapes corresponding to their genotype and experience condition. (Right) Individuals are colored by K-means clustering assignment, showing that the primary separation is SH from the rest of the conditions. Inset shows the % variance explained by the different PCs, with PC1 explaining the most variance. **(B)** Weights for each brain region for principal component (PC) 1 (left panel) and 2 (right panel) from the analysis in (A). The map regions (corresponding lateral coordinates) with strongest positive and negative values contribute most strongly to the variation between individuals. **(C)** As in B, weights for each brain region for PC1 (left panel) and 2 (right panel) are shown, in this case for PCA on data in which all cohorts were averaged for each condition. **(D)** Conditions projected onto PC1 show a separation of NH from the rest while PC2 axis shows a separation of SH from SW.

In the second PCA (Figure 7C, D), we sought to determine the major PNN density patterns that distinguish genotypes and conditions. Instead of averaging map numbers as in the previous analysis, we preserved data for each map number and averaged across the 5 cohorts for each condition and then performed PCA to determine the major distinguishing patterns. This analysis revealed patterns of PNN density that best distinguish NH vs. SH (PC1) and SW vs. SH (PC2) (Figure 7C, D). These PC patterns also reflect the medial/lateral and left/right asymmetries, thus confirming the anatomical and neurobiological distinctions in the previous figures. Overall, we observe that unsupervised analysis identifies these lateral/medial and left/right asymmetries as the major pattern that characterizes the differences in PNN distribution between experience and genotype.

### Individual mice exhibit strong laterality for PNN expression

As the previous data were an aggregate/average of five biological replicates, we were interested in determining if hemispheric biases in PNN density were seen in individual mice. For each mouse, we normalized PNN density of left hemisphere to the right hemisphere (Figure 8). In SS1 as a whole, a modest left hemisphere bias was seen in three out of five mice across conditions and genotypes (Figure 8A). Higher differences in left hemisphere bias is seen in subregions such as S1BF (Figure 8B) and S1ULp (Figure 8C). Interestingly, a decrease in the left hemisphere bias is observed in most of the SH mice, suggesting that SH brains have intact plasticity mechanisms that can be triggered by this social maternal experience to overcome the abnormal high PNN density lateralization in NH. Together, these results show that individual mice have differing hemispheric bias in PNN density in SS1, which may contribute, in a MECP2-dependent manner, to individual variability in responding to and consolidating new tasks involved in tactile sensation.

**Figure 8:**
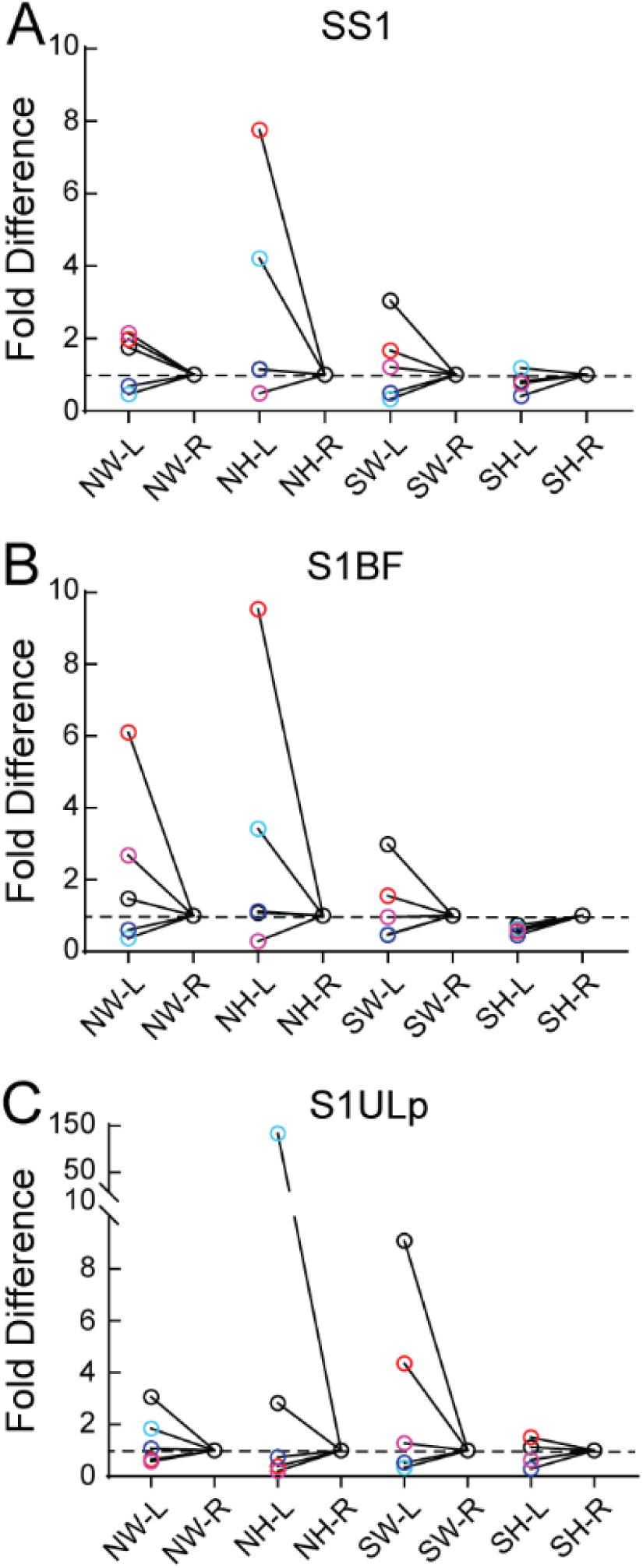
Individual mouse exhibits hemispheric bias for PNN density. **(A-C)** For each mouse, PNN density from the left hemisphere was normalized to the right hemisphere. This left-right hemisphere normalization revealed varying patterns of dominance. **(A)** In SS1, three NW mice exhibit left hemisphere bias of 2-fold, while two mice have a slight right bias (blue colors). After surrogacy, left hemisphere bias is increased to 4-fold in one mouse (black circle), while others remain similar to NW levels. Two NHs exhibit large fold differences favoring left hemisphere (red, blue circles). After surrogacy, dramatic loss in hemispheric bias is observed in the five mice brains. Similar trends with larger fold differences are seen in S1BF **(B)** and S1ULp **(C)**.

## Discussion

Given the long-standing and revitalized interest in extracellular matrix structures in the brain, we sought to systematically characterize high-intensity PNN expression in the whole SS1 in a model of adult experience-dependent plasticity, in relevant social behavioral conditions. To our knowledge, this is the first systematic characterization of PNN expression in the adult primary somatosensory cortex with nuanced information about subregions and laterality in individual mice.

In early postnatal cortical development, expression of PNNs increases progressively with the maturation of that network. An excellent example is the primary visual cortex where the developmental increase of PNNs is regulated by visual experience (Beurdeley et al., 2012; Hou et al., 2017). PNNs in mature primary visual cortex mainly surround the soma and proximal dendrites of PV+ interneurons (Celio, 1993; Hartig et al., 1992; Ueno et al., 2018). Mature PNNs are thought to be inhibitory for experience-dependent plasticity, as their increase in developing primary visual cortex correlates with the termination of the critical period and PNN removal in adult primary visual cortex restores plasticity, as measured by ocular dominance plasticity assays (Bavelier et al., 2010; Pizzorusso et al., 2002, 2006). These and studies in other brain regions (amygdala, hippocampus, piriform & auditory cortex) have suggested that PNNs are stable, long-term structures (Miyata and Kitagawa, 2017; Sorg et al., 2016; Ueno et al., 2019). Experiments involving the enzymatic removal of PNN by chondroitinase ABC (ChABC) or hyaluronidase injections in amygdala, hippocampus, piriform and auditory cortices have shown that synaptic plasticity can be reactivated (Banerjee et al., 2017; Gogolla et al., 2009; Kochlamazashvili et al., 2010; Krishnan, Lau et al., 2017; Pizzorusso et al., 2002; Thompson et al., 2018).

A word of caution: due to ease of immunostaining with WFA and manipulation experiments with ChABC, many studies now employ PNNs as markers for plasticity. Our characterization in adult brains shows that these structures are dynamic and have hemisphere- and subregion-specific expression, hinting at potential neural circuitry mechanisms involving laterality in mice. Systematic and careful analysis must be taken to fully characterize PNN expression in experimental design rather than using standard “representative” sample approaches in immunostaining.

What governs PNN dynamics in adults? Matrix metallopeptides and proteases are known to assist in remodeling extracellular matrix structures (Lorenzo Bozzelli et al., 2018; Lu et al., 2011; Miyata and Kitagawa, 2017). However, the contexts and mechanisms for inducing remodeling in adult brains are currently unclear. Some regional and temporal changes in PNN density have been described before (Ueno et al., 2018, 2019). However, systematic, finer scale whole-brain analysis of WFA expression across entire brain regions during development and adulthood has not been performed. Our study suggests PNNs, as measured by WFA immunostaining, may not be stable and static structures as once thought. In this study, we show that high-intensity PNNs of SS1 exhibit increased and decreased expression in a subregion-specific, hemisphere-specific manner, after maternal behavior experience. Currently, these differences in PNN expression between naïves and surrogates occur over one-two weeks (three – five days before pups are born plus six days of behavior before mice are perfused). The rate of PNN formation and remodeling, which might ultimately affect tactile perception and efficient pup retrieval, remains unknown.

In the Paxinos and Franklin atlas, anatomical subregions were classified based on structural connectivity studies. Based on these anatomical characterizations, we speculate that changes in PNN density in specific subregions could impact information processing. For example, when NW female mice learn maternal behavior to become surrogates, there is an increase in PNN density in the right S1BF, and a concomitant decrease in left S1FL (Figure 9), suggesting that these changes contribute to solidifying new synaptic contacts in S1BF, while promoting remodeling in S1FL. Together, these changes could ultimately help process new tactile information related to pups and the mother acquired by the whiskers and forelimbs. Furthermore, right hemisphere-specific PNN increases in S1FL and S1DZ suggests specific rewiring in subregions that could contribute to efficient information processing associated with laterality and dominant hemispheres (Figure 9). This idea of lateralization of function in the rodent brain has been a topic of longstanding yet of sporadic interest in the field (Glick and Ross, 1981; Kim et al., 2012; Soma et al., 2017).

**Figure 9:**
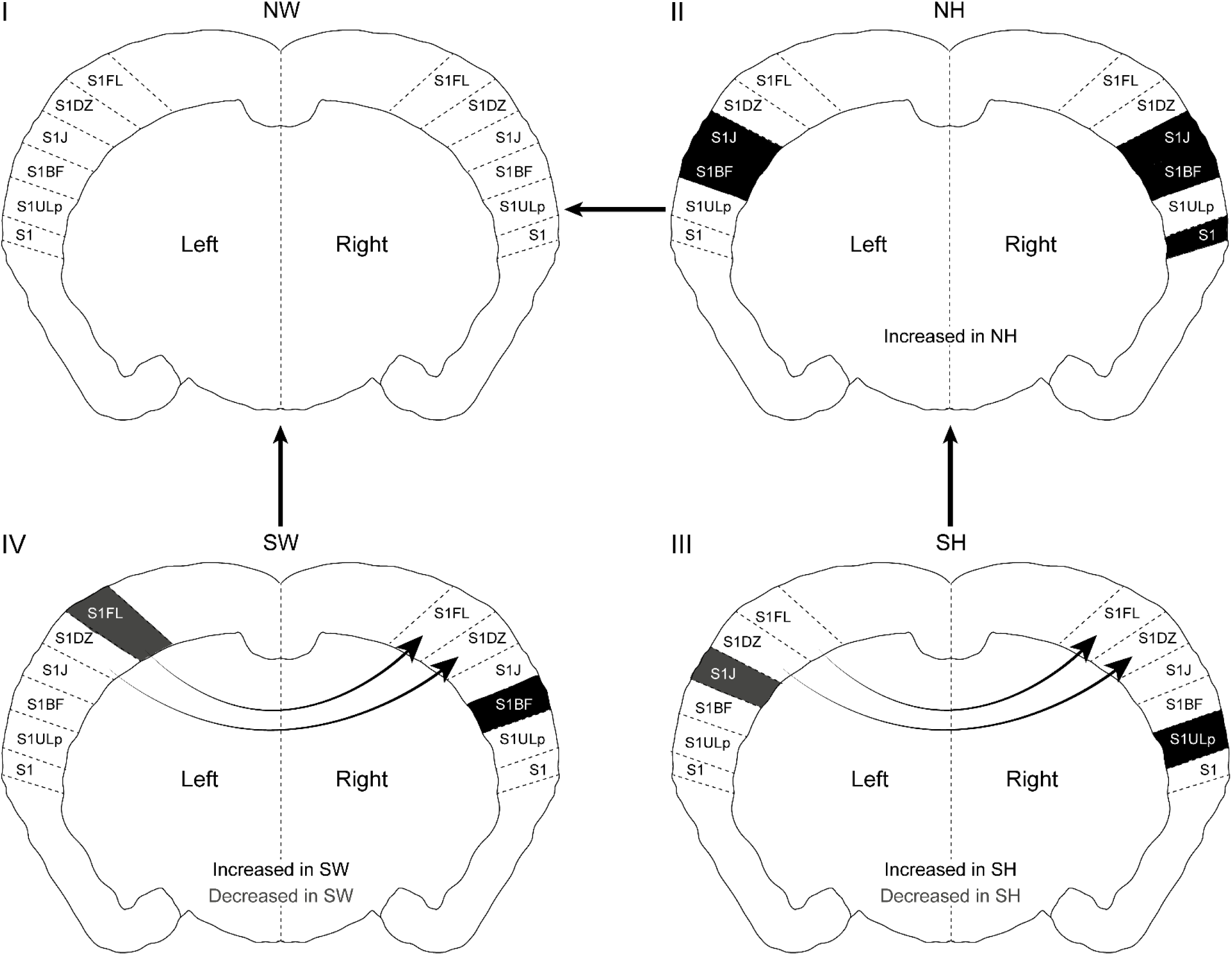
Summary of changes in PNN density between genotypes in maternal behavior context. (Quadrants I-IV) The changes in PNN density (grey and black shading) marked inside the brain slices denote comparisons between conditions connected by outside arrows between brain schemas. Arrows inside the brain schemas indicate hemispheric differences within genotype, with arrowheads pointing to the hemisphere with the higher PNN density. (IV→I) Comparing SW to NW, PNN density is increased in right S1BF and decreased in left S1FL regions of SW. Within SW, PNN density is increased in the right hemisphere, particularly in S1FL and S1DZ. Taken together, PNN density changes in these particular subregions could contribute to tactile perception in SW, ultimately leading to efficient pup retrieval. (II→I) NH had increased PNN density in specific subregions, compared to the NW, suggesting possible tactile perception issues before maternal experience, which could contribute to their inefficient pup retrieval performance. (III→II) SH has increased PNN density in the right S1ULp and decreased density in left S1J, compared to NH, suggesting possible compensatory plasticity mechanisms after maternal experience in Het. SH also displays increased right hemisphere PNN density increases in S1FL and S1DZ, similar to SW, suggesting that right hemisphere-specific increases in PNNs in S1FL and S1DZ might be important for processing tactile information during pup retrieval task.

Previously we showed that, in this adult female mouse model for Rett Syndrome (Het), PNNs were increased in a transient atypical manner in the auditory cortex, which correlated with their inefficient pup retrieval (Krishnan, Lau et al., 2017). Manipulating auditory cortex PNNs by ChABC injections or genetic reductions in Het significantly improved aspects of SH pup retrieval behavior, showing that PNNs play crucial roles in learning and executing this behavior (Krishnan, Lau et al., 2017). Current results suggest information flow, network activation and multisensory integration could be affected in Het, in specific cortical regions such as SS1 and auditory cortex (Krishnan, Lau et al., 2017; Morello et al., 2018). Further whole brain analysis on laterality in PNN density in the auditory cortex is warranted, especially due to reports suggesting left hemisphere-specific neural circuitry activation (Ehret et al., 1987; Marlin et al., 2015; Stiebler et al., 1997). Furthermore, MECP2 regulates tactile perception in SS1 of MECP2-deficient male mice (Orefice et al., 2016). However, it is currently unknown if PNNs are involved in these particular circuits and/or contribute to the observed tactile phenotypes.

The observed fine-scale changes in PNN expression in the adult primary somatosensory cortex before and after maternal behavior experience suggest specific hypotheses about connectivity and functional changes in subregions, specifically in barrel field and upper lip subregions of both hemispheres. *In vivo* electrophysiological and/or imaging studies, which measure dynamics of neural circuitry activation and processing in intact brains, would help prove these hypotheses.

## Author contributions

BYBL and KK supervised the project, designed experiments and analyzed data. KK, BYBL, DEL, ME and DGF performed the behavior experiments. BYBL, DEL, ME and BE performed sectioning and immunostaining. DEL imaged sections. BYBL, DEL, BE, KR, PS mapped sections to the reference atlas. BYBL, DEL, BE, ME, AK, PS, KR, AC, SHB and SR contributed significantly to manual counting of PNN structures. RPM performed and interpreted PCA analysis. DGF provided insightful and detailed comments on the manuscript. BYBL, DEL, RPM and KK wrote and edited the manuscript.

## Acknowledgements

We would like to thank the following students in helping us get started with this project and for technical contributions: Ashlee Tannehill, Taryn Lester, Ronald Dean Franz, Marty Lay and Simran Dayal. This work was supported by postdoctoral fellowship award to BYBL from RettSyndrome.org, research assistantships to DGF, BE, ME, PS from UTK and startup funds to KK from University of Tennessee-Knoxville.

## Abbreviations

PNN: Perineuronal nets
WFA: Wisteria Floribunda Agglutinin
PV+: Parvalbumin+ GABAergic neurons
MECP2: Methyl CpG-binding protein 2
NW: Naïve wild type
NH: Naïve *Mecp2* heterozygote
SW: Surrogate wild type
SH: Surrogate *Mecp2* heterozygote
PCA: Principal component analysis
SS1: Primary somatosensory cortex

**Subregions of SS1**
S1BF: barrel cortex
S1DZ: dysgranular zone
S1FL: Forelimb region
S1J: Jaw region
S1ULp: Upper lip
S1Tr: Trunk region
S1HL: Hind limb
S1: Undefined somatosensory cortex

